# Phosphorylation elicits bifurcated regulation of Na_V_1.5 by intracellular auxiliary proteins

**DOI:** 10.64898/2026.11.16.762840

**Authors:** Laurie Peverini, Iacopo Galleano, Lukas M. Fagerlund, Sitong He, Ruitao Jin, Janne Colding, Céline Marionneau, Ben Corry, Stephan A. Pless

## Abstract

The cardiac sodium channel Na_V1_.5 generates the upstroke of the cardiac action potential. Its inactivation is tightly controlled because shifts in steady-state inactivation (SSI), or incomplete pore closure generating a late current, are arrhythmogenic. Inactivation is modulated by tyrosine phosphorylation and protein-protein interactions (PPIs), but whether they intersect is unclear. Combining protein semi-synthesis, electrophysiology, biochemistry, fluorometry and molecular dynamics simulations, we demonstrate how phosphorylation at Y1495 in the DIII-DIV linker governs regulation by intracellular fibroblast growth factor homologous factors (iFGFs). We find that iFGFs act through two separable mechanisms: a depolarizing SSI shift that is switched off by phosphorylation, and a late current suppression that is independent of phosphorylation. Only the former requires an intact C-terminal domain/DIII-DIV linker interaction, and both differ between iFGF isoforms. Y1495 phosphorylation therefore does not abolish the iFGF interaction but redirects its output. This shows how a single post-translational modification selects between distinct PPI consequences.

## INTRODUCTION

The cardiac member of the voltage-gated sodium channel family, Na_V1_.5 (1), carries the fast sodium influx that drives the initial phase of the cardiac action potential. Its activity follows a complex gating cycle between closed, open, and physiologically important inactivated states (2). The sodium influx *via* Na_V1_.5 is terminated within milliseconds after channel activation by conformational changes underlying a process known as fast inactivation. Alternatively, sodium flux can also be reduced through a conformational change occurring from the closed state, known as steady-state inactivation (SSI) (3). The voltage-dependence of SSI determines the voltage range over which Na_V1_.5 is available for activation. Shifts in the voltage-dependence of SSI of only a few millivolts can therefore result in pathological conditions such as arrhythmias or sudden cardiac arrest (4,5). A related cause for Na_V1_.5-mediated pathogenicity is the development of late current due to defective or incomplete inactivation (6). This delays repolarization and can prolong the action potential, thereby dysregulating the Na^+^/Ca^2+^ homeostasis and disturbing the electrical conduction inside the heart (5). The voltage dependence and extent of Na_V1_.5 inactivation thus need to be precisely regulated to ensure the subsequent phases of the cardiac action potential.

Mechanistically, Na_V i_nactivation is driven primarily by the movement of voltage-sensing domains (VSDs) and by two key intracellular elements: the DIII-DIV linker, which carries the well-characterized Isoleucine-Phenylalanine-Methionine (IFM) inactivation particle that docks into its receptor site to allosterically stop sodium influx (7–12), and the C-terminal domain (CTD) whose contribution to inactivation is increasingly recognized (13,14). Interestingly, structural, functional, and computational studies show that the CTD directly contacts the DIII-DIV linker, and that this interaction is conserved across several mammalian Na_v s_ubtypes and appears dynamic and state dependent (14–19). This CTD/DIII-DIV linker interaction has thus emerged as an interface with potentially high relevance for the regulation of Na_v i_nactivation (20).

On the CTD side of this CTD/DIII-DIV linker interface, numerous protein-protein interactions (PPIs) have been described (21–23) that modulate Na_V1_.5 expression, sub-cellular localization and activity (24). Among those PPIs, intracellular fibroblast growth factor homologous factors (iFGFs, also known as FHFs (25,26)) interact with several voltage-gated sodium channel isoforms and regulate key functional properties, including maximum current (Imax), voltage-dependence of SSI (V_50S_SI) and late current development (27–32). The iFGF family comprises multiple splice isoforms that are differentially expressed across developmental stages, tissues, and pathological conditions (26,33–37). They bind to the EF-hand domain of the CTD (20,21,38), an interaction that likely also involves calmodulin (CaM) (39,40). The importance of iFGFs for cardiac physiology is further exemplified by the fact that iFGF variants are associated with cardiac disorders such as the Brugada syndrome (41) and ventricular tachycardia (42). Furthermore, variants affecting iFGF binding to the Na_V1_.5 CTD have likewise been associated with cardiac abnormalities (43).

In addition, the equilibrium between closed, open and inactivated states of Na_V1_.5 is also modulated by post-translational modifications such as phosphorylation (44,45). Proteomic approaches have identified more than 40 phosphorylation sites on Na_V1_.5 (46), with effects ranging from modulation of the expression to direct functional modulation (47,48). Interestingly, two serine phosphorylation sites within the Na_V1_.5 CTD have been identified as regulators of the Na_V1_.5-iFGF13 interaction (49). Phosphorylation at these sites reduces iFGF13 binding, which correlates with the loss of iFGF13-mediated effects on channel inactivation and late current reduction and highlights the important effect that phosphorylation can exert on PPI.

On the DIII-DIV linker side of the CTD/DIII-DIV linker interface, semi-synthetic protein engineering combined with electrophysiological recordings and molecular dynamic (MD) simulations have described how a single tyrosine phosphorylation site regulates Na_V1_.5 inactivation by impacting the binding of the nearby inactivation particle (50).

Thus, factors operating from either side of the CTD/DIII-DIV interface have been proposed to modulate inactivation: CTD binding partners (such as iFGFs) and at least one DIII-DIV linker phosphorylation site. But these have been only studied in isolation, leaving the question of a potential reciprocal crosstalk open. We therefore set out to directly assess if Y1495 phosphorylation, which profoundly alters inactivation (48,50), would also affect the functional impact of CTD PPIs, such as those involving iFGFs.

To this end we used a live-cell semi-synthetic method (tandem protein trans-splicing; tPTS (51)) paired with electrophysiology, biochemistry, voltage-clamp fluorometry (VCF) and molecular dynamic (MD) simulations to demonstrate an unexpected regulatory mechanism of iFGFs on Na_V1_.5 that is partially dependent on the phosphorylation state of Y1495 in the DIII-DIV linker. We further uncover the molecular basis for iFGF isoform-dependence of these effects and use MD simulations and pathogenic mutations to probe how Y1495 phosphorylation influences the dynamic interaction between the CTD and the DIII-DIV linker. Together, our data provide direct evidence for how phosphorylation can differentially impact Na_V1_.5 protein dynamics and selectively redirect the functional impact of CTD-binding PPIs.

## RESULTS

### Phos^1495^ abolishes iFGF A isoform-evoked SSI shift but spares late-current suppression

In light of the suggested interaction between the DIII-DIV linker and the CTD in Na_V c_hannels, we set out to investigate if functional effects elicited by CTD-binding proteins would be impacted by the phosphorylation state of the DIII-DIV linker. However, conventional approaches do not enable precise control over the phosphorylation state of defined sites in a cellular context: kinases/phosphatases or agents that activate/inhibit them affect all possible target sites and thus lack specificity. Site-directed mutagenesis can be used to ablate phosphorylation sites, but introducing phosphomimic side chains, such as glutamate, results in unwanted changes in size, charge and hydrogen bonding pattern (52). We therefore turned to a tPTS-based semi-synthetic approach to investigate if phosphorylation at Y1495, located in the α-helix upstream of the IFM particle in the Na_V1_.5 DIII-DIV linker, impacts the functional effects of PPIs on Na_V1_.5 (50,51). To generate Na_V1_.5 with an enzymatically stable phospho-Tyr mimic (a phosphonylated tyrosine analog) in position 1495, we recombinantly expressed the N- and C-terminal parts of Na_V1_.5 (amino acids (aa) 1 through 1471 (N) and 1503 through 2016 (C) of human Na_V1_.5) and injected a synthetic peptide X corresponding to part of the DIII-DIV linker (aa 1472 through 1502 of human Na_V1_.5), containing either a phosphonylated (non-hydrolysable phosphomimic) tyrosine (Phos^1495^) or a phenylalanine that cannot be modified by kinases (No Phos^1495^), into *Xenopus laevis* oocytes. (Fig. 1A) The N, C and X fragments are spliced together by two orthogonal split intein pairs, *Cfa*DnaE and *Ssp*DnaB^M86^ (50,51). In line with previous results, the removal of the Y1495 hydroxyl group in the No Phos^1495^ construct did not have a major functional impact compared with WT, while the Phos^1495^ led to a significant depolarizing shift in the voltage-dependence of SSI (V_50S_SI), presumably by destabilizing the IFM-bound state. This notion is supported by MD simulations from our previous study (50) (Fig 1B). The presence of the Phos^1495^ side chain also evokes a significant late current in two-electrode voltage clamp (TEVC) recordings (around 3%) that is not observed with No Phos^1495^ (Fig 1C).

**Figure 1.**
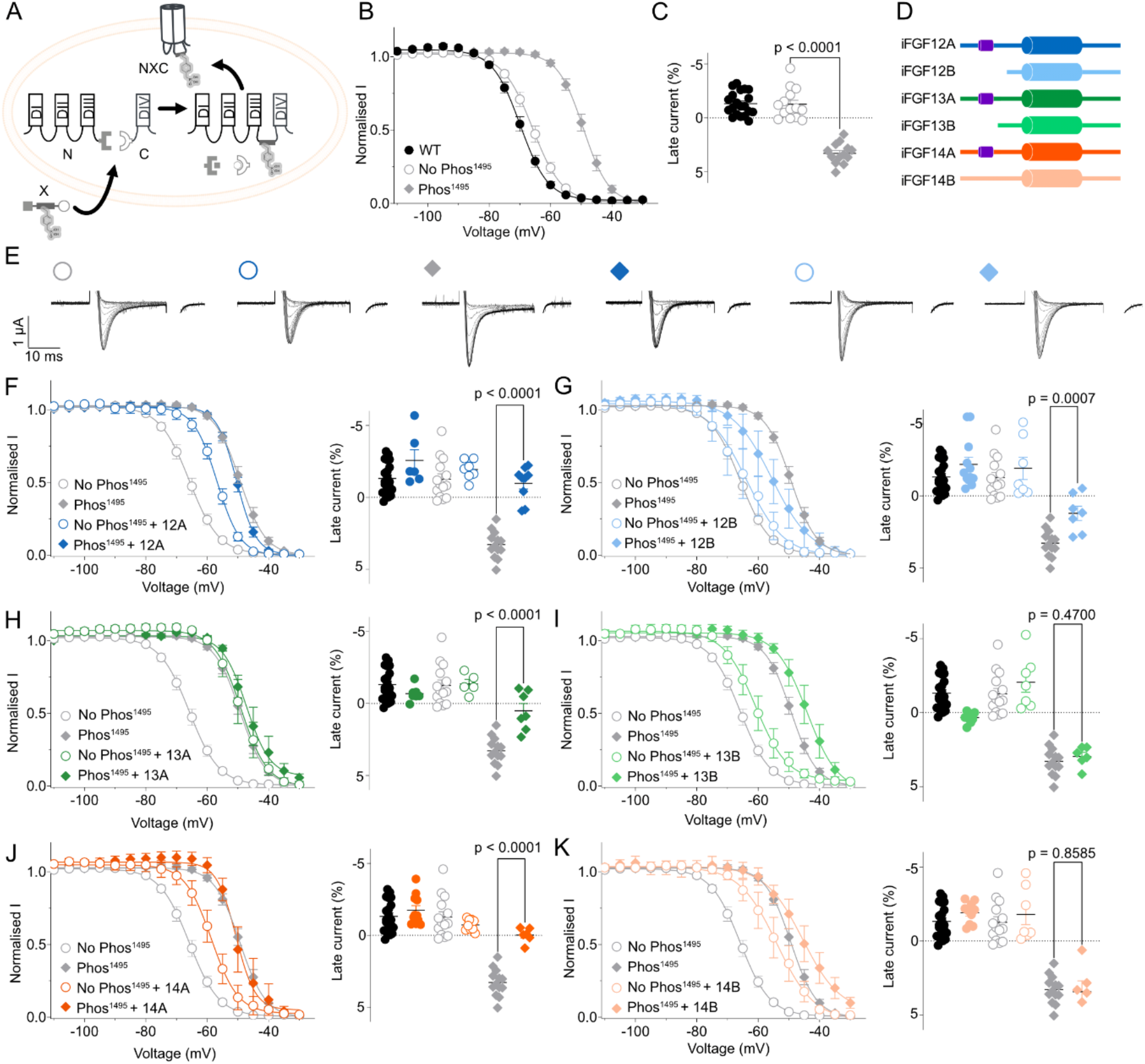
Phos^1495^ abolishes iFGF A isoform-evoked SSI shift but spares late-current suppression. **A.** tPTS strategy to express hNa_V1_.5 with and without non hydrolysable phosphomimic in Y1495. **B.** SSI curves for No Phos^1495^/Phos^1495^ compared to WT. **C.** Late current calculated for No Phos^1495^/Phos^1495^ compared to WT **D**. Schematic representation of the iFGF isoforms investigated in this study. **E.** Representative recordings of the conditions explored in panels F and G, corresponding to the 20 ms pulse of depolarizing voltage (-20 mV) part of the SSI protocol (see methods). **F-K.** left panel: SSI, right panel: late current for indicated iFGFs. Data shown as mean +/-SEM, with Welch’s t-tests for statistics (n=5-20).

We next examined the effects of iFGF co-expression on SSI of wild-type (WT) (Fig S1) and of Na_V1_.5-No Phos^1495^/Phos^1495^, with or without co-expression of six iFGF isoforms (iFGF12A, iFGF12B, iFGF13A, iFGF13B, iFGF14A and iFGF14B, Fig1D). TEVC recordings revealed differential effects of the studied iFGF isoforms co-expressed with WT Na_v1_.5 on V_50S_SI (Fig S1). We observed similarly pronounced shifts in V_50S_SI when co-expressing iFGF A types with No Phos^1495^ but, strikingly, no further shift on Phos^1495^ (Fig1 F, H, J, left panels). Smaller depolarizing shifts are observed with iFGF B subtypes (Fig1 G, I, K left panels), highlighting different functional effects between iFGF A and B isoforms on Na_V1_.5 inactivation. Importantly, the lack of iFGF12A/13A/14A-induced V_50S_SI shifts with the Phos^1495^ constructs is not due to saturation effects: Firstly, iFGF13B + Phos^1495^ and iFGF14B + Phos^1495^ are about 5 mV further right-shifted than Phos^1495^ alone. Secondly, we have shown previously that disease-causing Na_v1_.5 variants like Q1476R can induce more pronounced right-shifts in V_50S_SI than Phos^1495^ alone (50). Next, we investigated the effect of iFGFs on the late current for No Phos^1495^/Phos^1495^. We observed a significant reduction of the late current when co-expressing Phos^1495^ with all iFGF-A isoforms (Fig1 F, H, J, right panels). This reduction is also significant with iFGF12B, although to a lesser extent (Fig 1G, right panel), but not with iFGF13B and iFGF14B (Fig 1I, K, right panels).

To assess if the observed effects were specific to iFGFs, we also investigated the impact of an intracellular scaffolding protein, α-actinin-2, whose proposed interaction site resides within the Na_V1_.5 DIII-DIV linker (53), as well as CaM, which is proposed to interact with the CTD (39,40). Our experiments reveal that neither co-expression with α-actinin-2 nor overexpression or chelation of CaM (by co-expression of a peptide containing four consecutive IQ motifs, here termed Calmodulin chelator (CC)) affects V_50S_SI or late current in either No Phos^1495^ or Phos^1495^ (Fig S2). Our data suggest that one of the canonical iFGF effects (late current reduction) is independent of phosphorylation at Y1495, while the other (SSI shift) is abolished by phosphorylation at Y1495. This also implies that iFGFs still bind to Na_V1_.5 in the presence of Phos^1495^. To test this notion directly, we performed co-immunoprecipitation experiments between iFGF13A and full-length WT Na_V1_.5 or a mutant version carrying the Y1495E phosphomimetic mutant (Fig 2A). Strikingly, and in line with our finding that late current effects are preserved in Phos^1495^, we find the amount of co-immunoprecipitated iFGF13A to be indistinguishable between WT and Y1495E mutant (Fig 2B). This suggests that iFGF binding to the DIII-DIV linker is unaffected by phosphorylation at Y1495.

**Figure 2:**
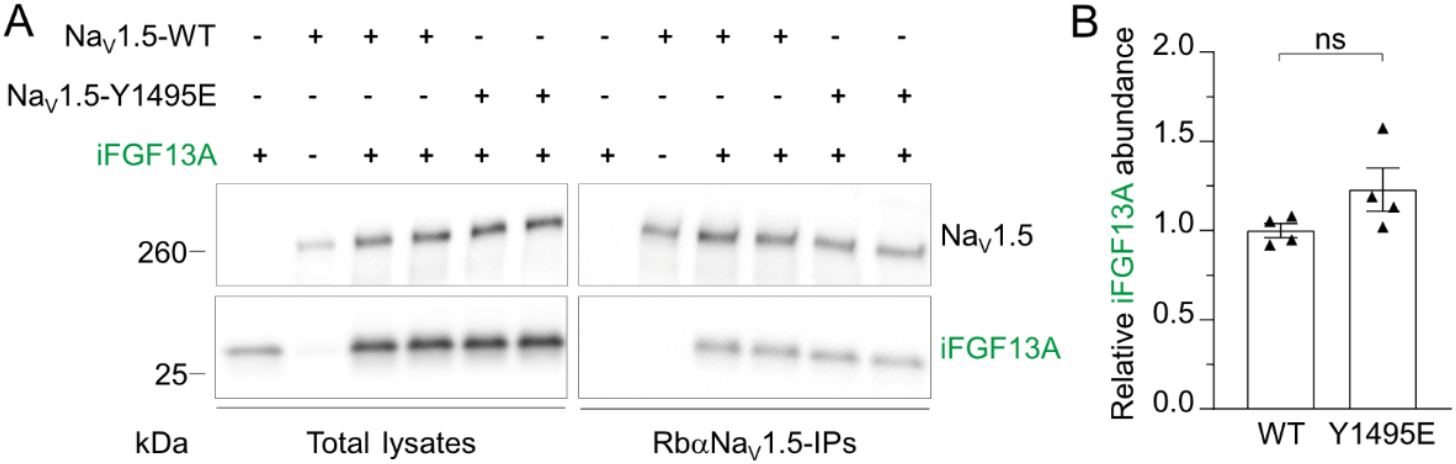
The direct interaction between iFGF13A and Na_V1_.5 is not affected by phosphorylation at Y1495. **A.** Representative western blots of the lysates (left panel) and the immunoprecipitates (right panel) with anti-Na_V1_.5 and anti-PAN-FHF-A antibodies. **B.** Relative mean ± SEM iFGF13A abundance in RbαNa_V1_.5-IPs from cells expressing Na_V1_.5-WT and Na_V1_.5-Y1495E. iFGF13A abundance in each IP were first normalized to immunoprecipitated Na_V1_.5, and iFGF13A abundance from Na_V1_.5-Y1495E-IPs were then expressed relative to abundances in Na_V1_.5-WT-IPs (Mann-Whitney test, p=0.1143, n=4 in each condition).

Together, these experiments reveal that Na_V1_.5 regulation by iFGF A isoforms is the consequence of two distinct mechanisms and that phosphorylation levels of Y1495 can differently redirect them without impairing binding to the DIII-DIV linker.

### Phos^1495^ impacts CTD/DIII-DIV distances in both inactivated and open states

To investigate how Y1495 phosphorylation impacts DIII-DIV linker dynamics during the Na_V1_.5 gating cycle, we performed triplicate 500 ns molecular dynamics (MD) simulations utilizing the full-length hNa_V1_.5 structures generated via ColabFold2. We modeled both the IFM-bound and unbound states, each with and without Y1495 phosphorylation. To select the starting structures, we generated 100 ColabFold2 models for each multiple sequence alignment (MSA) subsampling depth, ranging from 16 to full alignment. Finally, we clustered the resulting structures and selected the most representative model from each cluster yielding models of the IFM bound inactivated state and the IFM unbound partly open state (Fig 3 and Fig. S3A).

**Figure 3:**
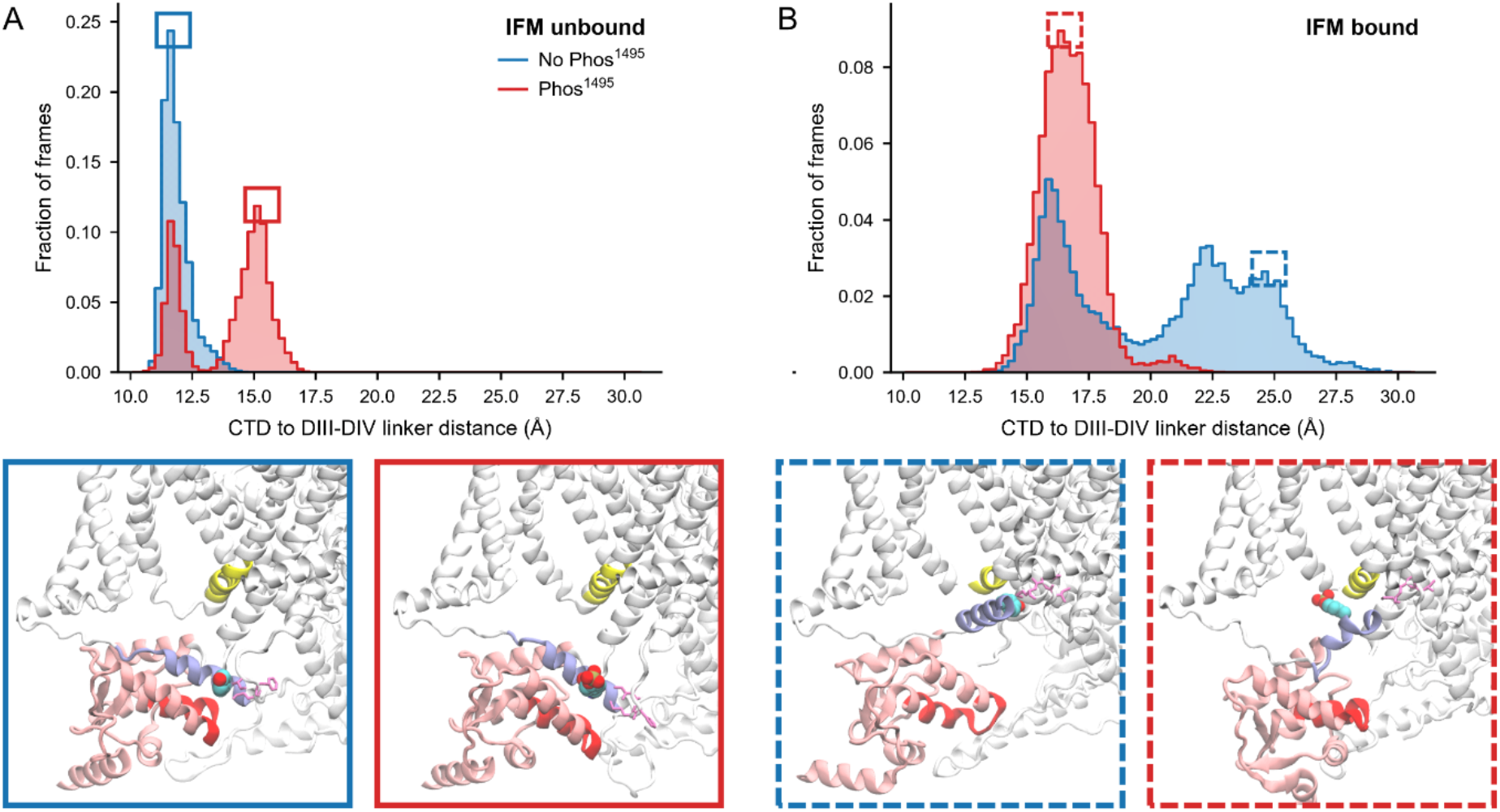
Phos^1495^ impacts CTD/DIII-DIV distances in both inactivated and open states. **A.** Fraction of frames at each COM distances between the Na_V1_.5 CTD and the DIII-DIV linker without (blue) and with (red) phosphorylation at Y1495 site from the IFM unbound (open) structure. Boxed insets below show structures corresponding to the conformations highlighted in the graph with the distance measured between the residues shown in blue and red ribbon. DIV S4-S5 linker is shown in yellow, DIII-DIV linker in blue, CTD in pink, Y1495 in balls and the IFM motif in pink sticks **B.** Same as A but for the IFM bound (inactivated) structure.

These simulations suggest that Y1495 phosphorylation repositions the DIII-DIV linker in a manner that depends on the conformational state. Backbone center-of-mass (COM) distances between the DIII-DIV linker (residues 1491–1507) and the CTD (residues 1854–1868) were monitored. From the IFM-unbound (inactivated) structure, the DIII-DIV linker/CTD separation was narrow and reproducible in the absence of phosphorylation (11.9 ± 0.6 Å) (Fig 3A blue). With phosphorylation at Y1495, the DIII-DIV linker slides relative to the CTD yielding a longer separation of approximately 15 Å in two replicates while the third remained at 11.8 Å, producing a pooled bimodal distribution despite the caveat of limited sampling (Fig 3A red). This suggests higher conformational plasticity that is conferred by Y1495 phosphorylation on the IFM unbound structure, with more conformations endowed with different DIII-DIV linker/CTD distances. By contrast, the simulations of the IFM-bound structure reveal longer distances without phosphorylation, with the DIII-DIV helix upstream IFM interacting more closely with the IFM binding site (Fig 3B), which likely recapitulates the more stable binding of the IFM particle in its site observed previously (50). Here, Phos^1495^ lowers the conformational plasticity of the IFM-bound structure perhaps by stabilizing a conformation with a DIII-DIV linker/CTD interaction partially disrupted or by stabilizing a partially inactivated state in which the IFM particle is less stably bound into its site. Comparing the DIII-DIV linker and the CTD residues contact patterns in the phosphorylated and non-phosphorylated systems, we find that dozens of residue pairs altered their occupancy by over 20 percentage points, with the majority of reproducible changes occurring downstream of CTD residue T1806 (84% IFM-unbound, 72% IFM-bound) (Fig. S3 B, C). Specifically, the unbound state showed a major reciprocal contact exchange at CTD residue F1855, while the bound state gained contacts between linker residue G1502 and CTD residues L1871 and M1875, aligning with reduced center-of-mass separation. Together, our MD work further emphasizes the critical effect of Phos^1495^ that appears to fundamentally modulate the conformational path of Na_V1_.5. When the IFM binds, it changes the location and environment of Y1495 substantially, which is also corroborated by voltage-clamp fluorometry (VCF) data revealing a robust quenching during depolarization with a fluorescent probe introduced at Y1495 position (Fig. S4). However, the position of Y1495 is substantially different when phosphorylated, which is in line with previous reports (50).

### CTD variant prevents iFGF-mediated SSI modulation only

To further probe the CTD-DIII/DIV linker interaction in the context of iFGF/Phos^1495^-induced modulation, we sought to destabilize it with a known pathogenic variant, E1784K, located in the CTD of hNa_V1_.5 (54– 56). This residue is proposed to interact directly with K1493 in the DIII-DIV linker *via* an electrostatic interaction. Changing the charge of either E1784 or K1493 promotes an hyperpolarizing shift of V_50S_SI, as well as the development of a late current (Supplementary Table 1) (20). We thus investigated the effect of the E1784K (EK) mutant expressed as No Phos^1495^ or Phos^1495^, with or without iFGF12A and iFGF13A. In contrast to WT, we found that the V_50S_SI shift induced by iFGF12A and iFGF13A is abolished for both No Phos^1495^ and Phos^1495^ on the EK mutant background (Fig 4B and C, left panels). We conclude that the iFGF-dependent V_50S_SI shift relies on an intact CTD/DIII-DIV linker interaction, which is disrupted by the EK mutant. Interestingly, the late current is significantly reduced when the Phos^1495^ EK variant is co-expressed with iFGF12A or iFGF13A (Fig 4B and C, right panels). This indicates that the iFGF-mediated effect on late current is not only independent of phosphorylation, but also independent of the destabilization of the CTD/DIII-DIV interaction, as promoted by the E1784K variant.

**Figure 4:**
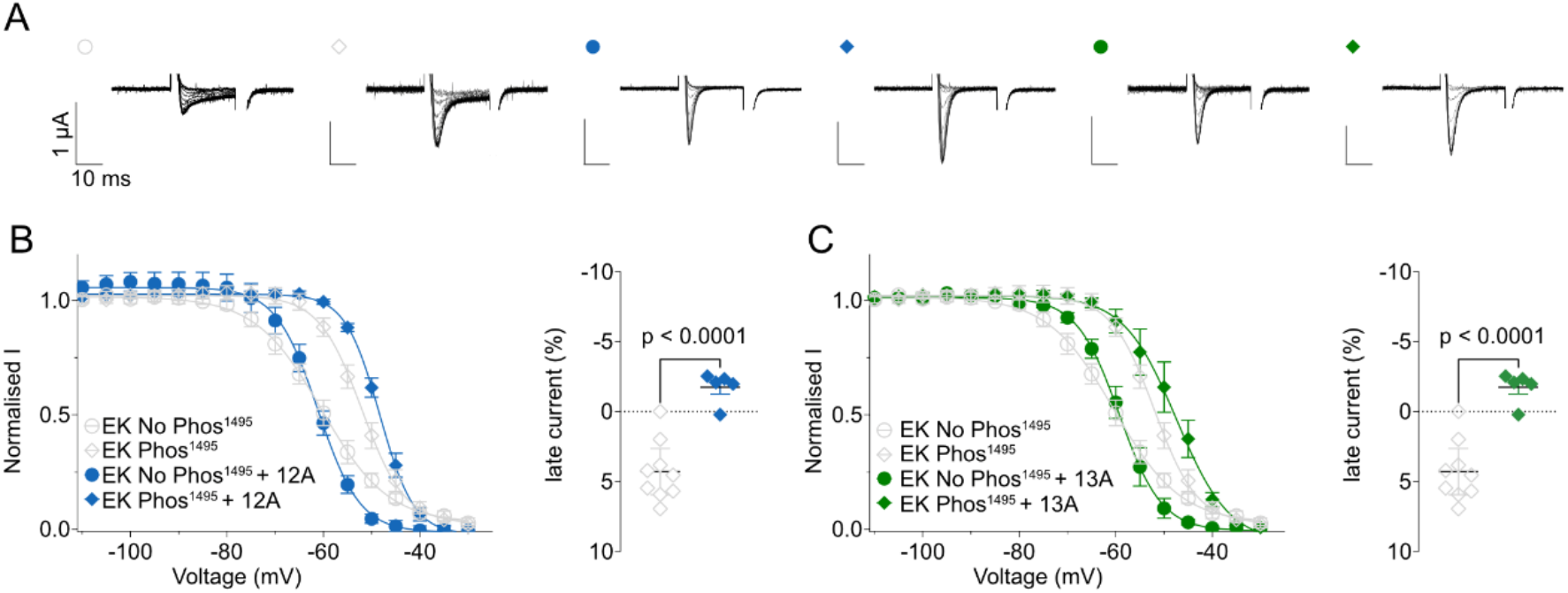
CTD pathogenic variant prevents iFGF-mediated SSI modulation but not late current reduction. **A.** Representative recordings of Na_V1_.5 No Phos^1495^/Phos^1495^ expressing E1784K mutant co-expressed with iFGF12A or iFGF13A. **B/C.** SSI curves and late current comparison for E1784K with iFGF12A (**B**) or iFGF13A (**C**). Data shown as mean +/-SEM, with Welch’s t-tests for statistics (n=5-12).

### The functional bifurcation is preserved with a Na_V1_.4 C-terminal domain

Since iFGFs are regulating numerous isoforms of Na_V,_ we set out to test if the effect of iFGF12A is conserved across Na_V c_hannels. Thus, we chose to investigate iFGF12A co-expression with a construct containing the CTD of Na_V1_.4. To this end, we took advantage of our tPTS approach to construct a chimera containing the DIV and CTD of Na_V1_.4 (corresponding to the C fragment), while the N and X fragments were derived from Na_V1_.5 (Fig 5A, see Fig S5 for sequence details). This chimeric construct showed robust expression and subtle differences in biophysical properties (Supplementary Table I). The voltage dependence of activation (V_50A_ct) for Na_V1_.5 (No Phos^1495^) was -38.1 +/- 0.4 mV, while V_50A_ct for the Na_V1_.5-Na_V1_.4 chimera was -30.8 +/- 1.0 mV. By contrast, V_50S_SI (No Phos^1495^) was similar for Na_V1_.5 and the Na_V1_.5-Na_V1_.4 chimera (Supplementary Table I). As observed above, the co-expression of the chimeric No Phos^1495^ construct with iFGF12A led to a significant depolarizing shift of V_50S_SI, but not with the chimeric Phos^1495^ construct. Interestingly, the depolarizing shift in SSI observed for the chimeric Phos^1495^ is somewhat greater than that for the Na_V1_.5 Phos^1495^ construct (approx.19 mV versus 16.5 mV). Further, the late current is reduced with the chimeric Phos^1495^ construct in the presence of iFGF12A, although to a slightly lesser degree: 0.8% of remaining late current with iFGF12A co-expressed with chimeric Phos^1495^, while there is no late current remaining with Na_V1_.5 Phos^1495^ upon iFGF12A co-expression. Together, these data suggest that despite subtle differences, the CTD of Na_V1_.5 and Na_V1_.4 behave largely similar with regards to the SSI shifts evoked by iFGF12A and phosphorylation at Y1495.

**Figure 5:**
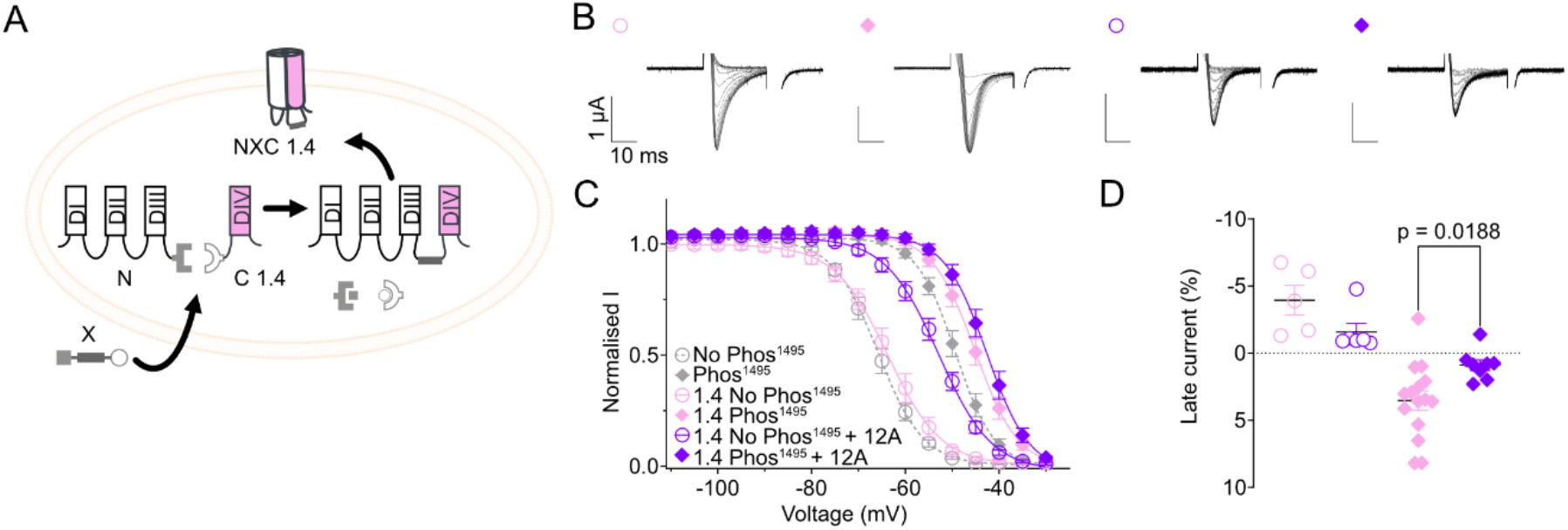
The functional bifurcation is preserved with a Na_V1_.4 C-terminal domain. **A.** tPTS strategy to generate Na_V1_.5-Na_V1_.4 chimera with Na_V1_.4 C part represented in pink. **B.** Representative recordings of Na_V1_.5-Na_V1_.4 chimera co-expressed (purple) or not with iFGF12A (pink) and comparison with Na_V1_.5 tPTS construct (grey, dashed lines) **C.** SSI curves **D.** Late current. Data shown as mean +/-SEM, with Welch’s t-tests for statistics (n=5-20).

### iFGF N-terminal domains are not sufficient for late-current suppression

Previous studies showed that the core domain of iFGFs is the principal interacting point with the CTD of Na_V1_.5 (39). Additionally, our results and previous reports highlight that the B isoforms, which possess a shorter or altered NTD, appear less efficient at reducing late current (see Fig 1 and (29)). Since the NTDs are the most divergent domain between A and B isoforms, we hypothesized that the NTDs of the A isoforms could explain the differences observed between iFGF A and B isoforms. We therefore decided to investigate the effects of isolated iFGF N-terminal domains (NTD) of two iFGF isoforms, iFGF12A and iFGF13A. We designed constructs containing the N-terminal residues that are different between iFGF12A and iFGF12B (amino acids 1-66 of iFGF12A, referred as NTD12A); and iFGF13A and iFGF13B (amino acids 1-62 of iFGF13A, referred as NTD13A) (Fig 6A and Fig S6). We then co-expressed these constructs with No Phos^1495^ and Phos^1495^. Co-expression of No Phos^1495^ with NTD12A reveals a depolarizing shift of V_50S_SI that is similar to that of full-length iFGF12A (9 mV for iFGF12A vs. 7 mV for NTD12A). We observed a lesser shift with NTD13A vs iFGF13A (17 mV for iFGF13A vs 8 mV shift with NTD13A). Surprisingly, we observed no reduction in late current when Phos^1495^ was co-expressed with NTD13A, whereas the late current was significantly reduced upon NTD12A co-expression. Interestingly with iFGF12B (whose sequence is closely related to iFGF12A without NTD12A), the late current is similarly reduced (see Fig 1 and Supplementary Table 1). Since the late current is not reduced by the co-expression of NTD13 (Fig 6D) or by the co-expression with iFGF13B (Fig 1I), full-length iFGF13A appears to be necessary to observe the reduction of the late current. This is different for V_50S_SI, where a change in the voltage dependence is observed, although this is less pronounced for the iFGF NTD constructs. On the contrary, iFGF12B and NTD12A both significantly reduced the late current, although to a lesser degree than iFGF12A.

**Figure 6.**
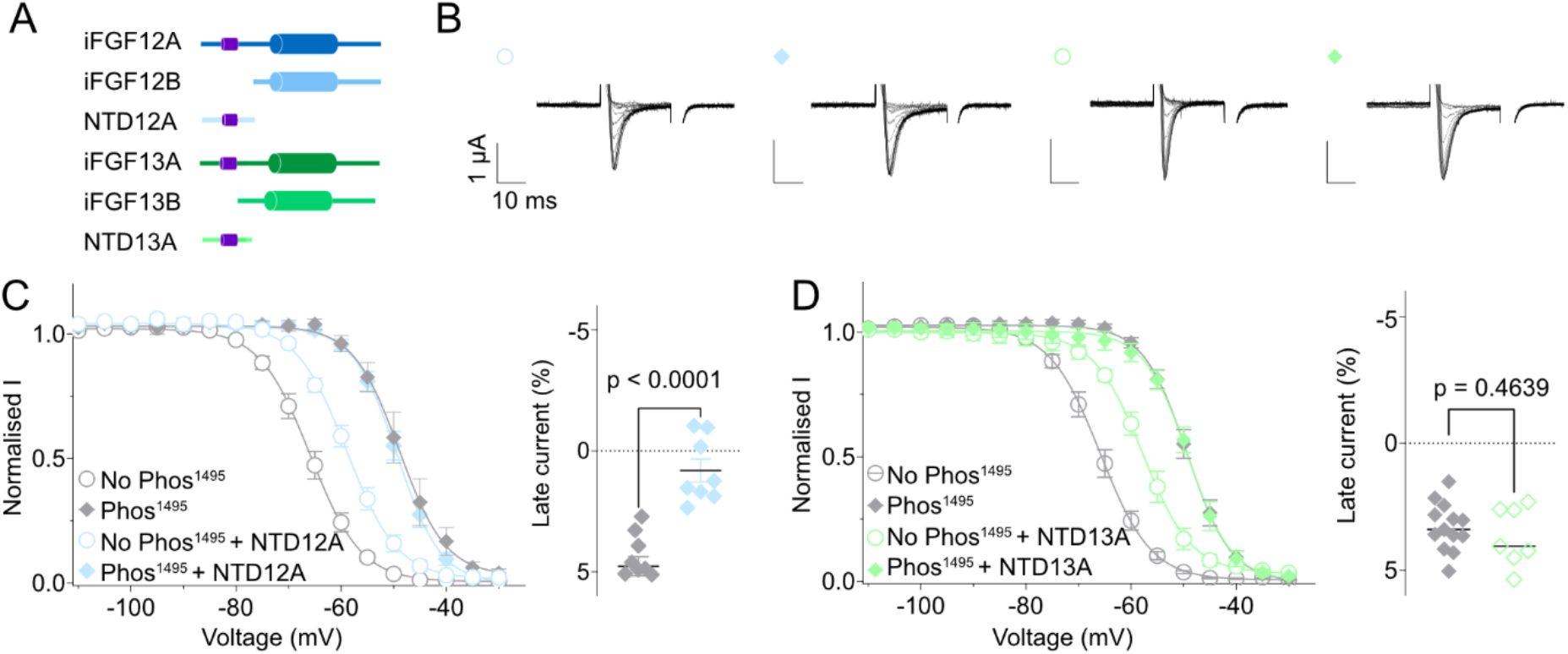
iFGF N-terminal domains are necessary but not sufficient for late-current suppression. **A.** Schematic representations of constructs NTD12A and NTD13A constructs co-expressed in the experiments compared to their corresponding iFGFs isoforms. **B.** Representative recordings of No Phos^1495^ and Phos^1495^ co-expressed with and without NTD12A and NTD13A **C/D.** NTD12A (**C**) and NTD13A (**D)** SSI curves and late current properties. Data shown as mean +/-SEM, with Welch’s t-tests for statistics (n=5-20).

Overall, the iFGF NTDs appear to play an important role in late current reduction; and the differences between NTD12A and NTD13A co-expressed with Phos^1495^ further highlight the subtype-dependent way in which iFGFs modulate Na_V1_.5 function.

## DISCUSSION

Here we combined live-cell protein semi-synthesis with electrophysiology, co-immunoprecipitation, VCF and MD simulations to establish how a single phosphorylation site in the Na_V1_.5 DIII-DIV linker governs channel regulation by intracellular auxiliary proteins. Tandem protein trans-splicing allowed us to compare Na_V1_.5 channel populations that were either unphosphorylated or fully phosphorylated at the Y1495 site and revealed that iFGF A isoforms act on Na_V1_.5 through two distinct mechanisms: a depolarizing shift in V_50S_SI that is switched off by Phos^1495^, and a reduction of Phos^1495^-induced late current that occurs irrespective of the phosphorylation state of Y1495. MD and VCF show that Y1495 resides in a highly mobile environment and that its phosphorylation reshapes the CTD/DIII-DIV linker interaction in a state-dependent manner, while the pathogenic CTD variant E1784K uncouples the CTD/DIII-DIV linker interaction and selectively removes the SSI shift elicited by iFGF modulation. Both effects are preserved in a Na_V1_.5-Na_V1_.4 CTD chimera, yet they differ markedly between iFGF splice isoforms. Phosphorylation at Y1495 thus acts as a molecular bifurcation that determines which of the two iFGF-mediated mechanisms is available to the channel.

### Phosphorylation reshapes the CTD/DIII-DIV linker interaction

We used MD simulations to investigate the mobility of the DIII-DIV linker in a full-length channel context. In the IFM-unbound (open) model, Phos^1495^ increased the center-of-mass distance between the CTD and the DIII-DIV linker and permitted sampling of multiple conformations, whereas the non-phosphorylated channel occupied mainly a single conformation. In the IFM-bound (inactivated) model the relationship was inverted, with the larger distances observed in the absence of phosphorylation, consistent with more stable IFM engagement (50). Phosphorylation further redistributed the contact interface, predominantly downstream of residue 1806 in the IFM-bound state and upstream of it in the IFM-unbound state. Y1495 phosphorylation therefore does not simply weaken the CTD/DIII-DIV linker interaction but rather re-routes it in a state-dependent manner, thereby profoundly impacting the gating cycle of Na_V1_.5, in line with reports showing this interface as highly dynamic across conformational states in other Na_V s_ubtypes (17).

### Phos^1495^ and E1784K destabilize the CTD/DIII-DIV linker interface differently

Destabilization of the CTD/DIII-DIV linker interface can be elicited by the Brugada/LQT3 E1784K variant that disrupts the electrostatic contact between E1784 in the CTD and K1493 in the DIII-DIV linker. Both E1784K and the charge-reversal K1493E produce a hyperpolarizing shift in V_50S_SI together with a pronounced late current (20,54). The presence of Phos^1495^ likewise generates a late current but shifts V_50S_SI in the opposite (depolarizing) direction, therefore presumably perturbing the CTD/DIII-DIV linker interaction through a mechanism distinct from that of E1784K.

Testing iFGFs against this E1784K-induced destabilization of the CTD/DIII-DIV linker interface revealed again a separation of the two iFGF-related mechanisms on Na_V1_.5 activity. On the E1784K background, neither iFGF12A nor iFGF13A shifted V_50S_SI in the No Phos^1495^ or Phos^1495^ conditions, while the reduction of Phos^1495^-evoked late current was fully retained. The SSI shift elicited by iFGF modulation thus requires an intact CTD/DIII-DIV linker interaction, which either Phos^1495^ or E1784K are sufficient to disable. By contrast, the late-current reduction does not, although iFGFs remain bound under both conditions.

### Functional impact of iFGFs is conserved among Na_V c_hannels

We next sought to determine whether the effects observed with Phos^1495^ and iFGFs are dependent on the exact CTD sequence context by testing it against a Na_V1_.5/Na_V1_.4 CTD chimera. This chimera displays 78% sequence similarity in the C part of the intein construct, with most differences occurring towards the end of the CTD. The effects evoked by iFGF12A on V_50S_SI and late current were notably similar for the Na_V1_.5 and Na_V1_.4 CTDs, suggesting a mechanism of modulation by FGFs that extends beyond just the Na_V1_.5 subtype. We did, however, observe subtle differences in the extent of V_50S_SI shift and the extent of reduction in late current, which underlines the importance of the exact CTD sequence context for subtype-specific biophysical properties.

Additionally, this experimental approach highlights tPTS as a convenient way to build chimeric constructs through a ‘cut and replace’ process. We anticipate this to prove useful in other contexts that require *in cellulo* chimera generation.

### Differential impact by iFGF parts and isoforms

Finally, we describe that iFGF A isoforms appear to be required to fully reduce the late current promoted by Phos^1495^, while iFGF B isoforms or just the NTDs do not, or do so to a lesser extent. These experiments highlight the highly variable roles of the different iFGF isoforms (29,31,35,57). Comparing the effects of NTD12A and NTD13A also reveals that only a few residues in the NTD of iFGFs have pronounced effects on their action on Na_V1_.5. NTD12A and NTD13A share high sequence similarity, with their N-terminal parts being identical for the first 18 and last 5 last residues. We therefore hypothesize that the determinants of the differential effect are located within the intervening areas. Since there are numerous splice isoforms of iFGFs expressed in excitable cells (32,33), their relative expression level and the splice regulation could be a way of fine-tuning Na_Vs_ function in different conditions (26,29). Future studies that follow the expression and splicing pattern of the different iFGF isoforms in different cellular or pathological contexts will therefore be of great interest to understand Na_V m_odulation by iFGFs.

### Implications for cardiac physiology and beyond

Under physiological conditions, both iFGF expression levels and the extent of tyrosine phosphorylation at Y1495 are dynamic and both are upregulated in stress or pathological conditions. Case in point, iFGF12A expression is reported to be increased in heart failure and iFGF13 is upregulated in cardiomyocytes under pressure overload conditions (58–60). Similarly, Y1495 is phosphorylated by Fyn kinases (45,48) and kinase activity is often increased under pathological conditions or stress, as underlined by β-adrenergic simulation triggering phosphorylation of cardiac ion channels (61). Together with our data showing an increased window current with both Y1495 phosphorylation and most iFGFs (*via* the depolarizing V_50S_SI shift), this suggests several routes towards an increased Na_V1_.5 activity in pathologic contexts.

We also study the impact of iFGF14 isoforms, whose expression has mainly been probed in the brain, even if the presence of Na_V1_.5 in the brain is still controversial (4,62). Interestingly, we show that iFGF14A impacts Na_V1_.5 function similar to what is seen with iFGF12A and 13A. This is relevant because we show that iFGF regulation, including its dual modulation by phosphorylation, likely extends to other Na_V s_ubtypes, which points to our data having uncovered regulatory mechanisms that apply beyond Na_V1_.5 regulation by iFGF12A and 13A only. It is also noteworthy that Na_V1_.5 is expressed in the gastro intestinal tract (4) and in cancer cells (63,64), and that iFGF12 is overexpressed in some cancers (34).

Together, we demonstrate that iFGFs tune Na_V1_.5 activity through inducing phosphorylation-dependent shifts in V_50S_SI or late current reduction, thus enabling a precise calibration of Na_V1_.5 function under different physiological and pathological conditions.

### Potential limitations

Our work underlines that tPTS represents a powerful approach to study the impact of a specific phosphorylation site and we demonstrate that this method is further an efficient way of generating and studying chimeric constructs. The approach has, however, some limitations. First, to allow direct RNA (N and C fragments) and peptide X injection we use *Xenopus laevis* oocytes, which have a cellular environment different from e.g. cardiomyocytes. Secondly, we only compare Na_V1_.5 populations with either 0% or 100% phosphorylation. Additionally, it is important to highlight that there has historically been considerable variability in electrophysiological data reported for Na_V1_.5 (65), which is why we primarily focus on comparing relative differences observed between the different conditions and apply strict exclusion criteria in our dataset (see Methods). It is also important to consider that we only investigate a single Na_V1_.5 phosphorylation site in this study, while recent phospho-proteomics in mouse ventricles has discovered at least 42 more (46) and some of them may impact iFGF13 binding via different mechanisms (49). Furthermore, iFGF13 itself has been reported to harbor at least nine phosphorylation sites, raising the possibility that phosphorylation dynamically regulates both partners of the Na_V1_.5-iFGF complex (66). This suggests a more complex situation in living organisms that motivates experimental approaches beyond single-site studies in the future.

### Conclusion and Outlook

Together, these findings establish phosphorylation at Y1495 site as a switch that selects which of two iFGF-mediated mechanisms is available to modulate Na_V1_.5 function, showing that a single post-translational modification can redirect the output of a protein-protein interaction without abolishing it. Since both channel phosphorylation and iFGF isoform abundance change in cardiac disease, this balance is likely remodeled when inactivation defects become arrhythmogenic. Correcting such defects by targeting phosphorylation sites or auxiliary-protein interfaces, rather than the channel pore, may thus represent a promising therapeutic strategy.

## MATERIAL AND METHODS

### Chemicals

Chemicals and salts were purchased from Sigma-Aldrich, VWR Chemicals, ThermoFischer Scientific, Acros Organics and Panreac Applichem.

### Constructs and mutants

Human Na_V1_.5 (canonical sequence – Uniprot reference Q14524-1, codon-optimized), the inteins constructs design corresponding to N- and C-parts of Na_V1_.5 have been previously described (50,51).. All iFGFs (canonical sequence, codon-optimized, Uniprot references P61328-1 for iFGF12A, P61328-2 for iFGF12B, Q92913-1 for iFGF13A, Q92913-2 for iFGF13B, Q92915-1 for iFGF14A and Q92915-2 for iFGF14B) have been bought as expression-ready plasmids in pUNIV (plasmid #24705 addgene) from Twist Biosciences. Variants studied are introduced by site-directed mutagenesis using Q5 High-Fidelity DNA polymerase (Qiagen). Correct modifications are assessed by whole-plasmid sequencing (Eurofins genomics). RNA for oocyte injection is produced after linearization (NotI restriction enzyme) by T7 transcription kit (mMESSAGE mMACHINE™ T7 Transcription Kit, ThermoFischer Scientific).

### Peptide synthesis

Peptides X (with phosphonylated residue in Y1495 or Y1495F) were synthesized in a previous study by solid-phase peptide synthesis (51). Note that the residue N1472 is mutated into a cysteine to ensure the splicing efficiency and that the mutation K1479R is added. The impact of these mutations have already been assessed in our previous study (51).

### Expression in *Xenopus laevis* oocytes

Oocytes lobes from *Xenopus laevis* are surgically removed from the animal (anaesthetized in 0.3% tricaine, under license 2014-15-0201-00031, approved by the Danish Veterinary and Food Administration). The ovarian lobes are digested with collagenase type I (0.5 mg/mL) in OR2 buffer (in mM: 82,5 NaCl, 2,5 KCl, 1 MgCl_2_, 5 HEPES, pH 7.4). Stage V-VI oocytes are injected (nanoliter 2010 micromanipulator, WPI) with 30 nL of a mixed solution containing cRNA encoding the N- and C-inteins constructs (250ng/nL). After 24 hours of incubation in ND96 complemented with antibiotics (in mM: 96 NaCl, 2 KCl, 1 MgCl_2_, 1.8 CaCl_2_, 5 HEPES, supplemented with 0.275 g/L sodium pyruvate, 0.09 g/L theophylline, 0.05g/L gentamycin and 0.05 g/L tetracycline pH 7.4), 20 nL of peptide is injected (250 μM dissolved in Milli-Q water). The oocytes are recorded between 24 and 48 hours of expression (incubation at 18 degrees).

### Two-electrode voltage clamp recordings and voltage clamp fluorometry recordings

Voltage-clamp sodium currents are recorded with a two-electrode voltage clamp amplifier (OC-725C, Warner Instruments) associated with an Axon digidata 1550 (Molecular Devices) and pClamp10 software (Molecular Devices). Oocytes are impaled with glass microelectrodes with resistances between 0,1 and 0,8 MΩ filled with 3M KCl solution. Oocytes are clamped at -100 mV if not otherwise stated. ND96 is used as recording buffers: (ND96 in mM: 96 NaCl, 2 KCl, 1 MgCl_2,_ 1.8 CaCl_2,_ 5 HEPES, pH 7.4) and constantly perfused. For VCF experiments, oocytes are clamped in a custom-made chamber with their animal pole towards a water-immersed objective (IX73 with LUMPlanFL N 40x, Olympus) of an inverted microscope. A halogen lamp (Olympus TH4-200) is used for excitation with a filter set containing 377/50 nm excitation filter, a 409 nm dichroic mirror and a 490/60 nm emission filter (Chroma). The emission is detected *via* an avalanche photodiode (APD, Photomax200, Dagan Corporation) that subtracts baseline fluorescence. During VCF recordings, between 10 and 50 repeats of the same protocols are collected for the same signal and are averaged automatically in pClamp10.7.

### Protocols

To determine V_50A_ct, oocytes are clamped and held at -100 mV, and sodium currents are measured at the peak for voltage steps of 50 ms ranging from -80 mV to +40 mV (in 5 mV increments). To determine V_50S_SI, a 500 ms pre-pulse from -100 mV to -20 mV (in 5 mV increments) is followed by a 20 ms voltage step to -20 mV. Late current is calculated during SSI protocol for the recording of the pulse at -30 mV and represents the mean current between 400 ms and 420 ms after the peak current and is expressed as a percentage of the maximal current.

### Data analysis

Data considered have passed several exclusion criteria to ensure reproducibility. We considered recordings displaying minimal currents of -0.5 μA (except for one construct – see Supplementary Table 1) and where maximal current is maximum 3 μA for activation protocol to ensure a correct clamp, maximal leak considered was -0.1 μA. At least two batches were recorded for each construct. Number of repeats (n) are displayed in Supplementary Table 1. Raw data analysis was conducted in Excel and prismX (GraphPad). The data were fitted to Boltzmann sigmoidal curves to obtain the values of V_50A_ct and V_50S_SI. For VCF recordings, we did not analyze current-related parameters because the currents were too high to ensure a fast clamp, thus we focused on fluorescence properties. In VCF recordings figure legend, ΔF/F is an apparent ΔF/F calculated versus the baseline fluorescence in the beginning of the recording because fluorescence acquisition is normalized to zero prior to shown recordings. For VCF delay calculations, peak I and peak F are extracted in the same period, between 100 and 105 ms. The kinetic calculations are performed by mono-exponential fitting of fixed period in the fluorescence signal, 6 ms for tau_on a_nd 20 ms for tau_off._ Statistical tests used are Welch’s t test if not otherwise stated.

### Molecular dynamic simulations

Full length model: To generate full length starting conformations of Na_V1_.5 we generated 100 models using ColabFold with a full multiple sequence alignment (generated with colabfold_search) and with multiple sequence alignment subsampling ranging from 16 to 5120 to generate additional conformational diversity (67). The resulting models were clustered with a cutoff of 1.5 Å to yield 14 structural clusters in total with the top two representing 36% and 24% of models in total. The first of these had the IFM motif bound to its receptor and a closed activation gate, while the second had IFM unbound and a partially open activation gate in line with the diversity seen in previous models of the voltage gated channel family (17). Calmodulin was modelled onto the channel CTD with MODELLER 10.4 (68), using the Nav1.5 CTD–Ca²⁺/CaM interface of the 4JQ0 ternary structure as a template (69).

System construction: All simulation systems were built with the CHARMM-GUI Membrane Builder (70) and described by the CHARMM36m all-atom force field for protein (71), CHARMM36 for lipids, and the CHARMM-modified TIP3P water model. The two most populated model clusters described above (IFM-bound and IFM-unbound) were taken forward as starting structures with disordered intracellular loops removed. each with either unmodified tyrosine or phosphotyrosine. The native N-terminus (Met1) was retained as a standard charged amine; the N-termini created at the internal chain breaks and at the CaM terminus were acetylated, and all C-termini were capped as N-methylamide. All histidine residues were modelled as the neutral Nδ-H tautomer, five disulfide bonds were retained, and four Ca²⁺ ions were bound to CaM. Each complex was embedded in a 1,2-dioleoyl-sn-glycero-3-phosphocholine (DOPC) bilayer and solvated with explicit water together with Na⁺ and Cl⁻ ions to a bulk concentration of 0.15 M, with the Na⁺ excess set to render each system electrically neutral. The resulting systems contained 407,821–417,823 atoms in periodic boxes of approximately 14.3 × 14.3 × 19.1–20.1 nm.

Simulation parameters: Simulations were performed using the GPU-accelerated GROMACS software package (version 2023) (72). The LINCS algorithm (73) was applied for resetting constraints on covalent bonds to hydrogen atoms, which allowed 2 fs time steps for MD integration during production. The particle-mesh Ewald algorithm (74) was used for calculating electrostatic interactions, with a real-space cut-off of 12 Å, and the Verlet grid cut-off-scheme (75) was applied for neighbour searching, using an update frequency of 20 and a cut-off distance of 12 Å for short-range neighbours. A 12 Å cut-off was applied to account for van der Waals interactions, using a smooth force-switching function starting at 10 Å. Periodic boundary conditions were utilized in all directions. Solute (protein and Ca²⁺), membrane and solvent groups were treated independently for temperature and pressure coupling to increase accuracy, and centre-of-mass motion was removed every 100 steps.

Minimisation and equilibration: Equilibration utilized steepest descent energy minimization followed by eleven sequential steps of equilibration (two of 100 ps in the NVT ensemble and nine of 0.1–2 ns in the NPT ensemble, approximately 16.3 ns in total, using a 1 fs time step for the first three steps and 2 fs thereafter) with a gradual decrease in the restraining force applied to protein and lipid atoms. Over the first six steps the lipid position and dihedral restraints were released while the protein was held near its initial conformation, allowing the bilayer to pack around the channel; over the final five steps the protein backbone and side-chain restraints were released progressively. During the equilibration stages, the temperature was maintained at 303.15 K using a velocity-rescaling thermostat (76) with a time constant of 1.0 ps. From the third step onward, the pressure was maintained at 1.0 bar by semi-isotropic application of a stochastic cell-rescaling barostat (77), with a time constant of 5.0 ps and compressibility of 4.5 × 10⁻⁵ bar⁻¹.

Production simulations: During production of molecular dynamics, the temperature was maintained at 303.15 K using a Nosé–Hoover thermostat (78) with a time constant of 1.0 ps, and the pressure was maintained at 1.0 bar using the Parrinello–Rahman barostat semi-isotropically (79), with a time constant of 5.0 ps and compressibility of 4.5 × 10⁻⁵ bar⁻¹. Three replicas each lasting 500 ns were run for each of the four simulation systems, giving twelve trajectories and 6 µs of aggregate sampling, with coordinates saved every 100 ps.

### Culture and Transient Transfections

Human Embryonic Kidney 293 (HEK-293) cells were maintained in Dulbecco’s Modified Eagle’s Medium (DMEM, Life Technologies) supplemented with 10% fetal bovine serum, 100 U/ml penicillin and 100 μg/ml streptomycin, in 37°C, 5% CO_2_: 95% air incubator. Cells were transiently transfected at 60-70% confluence in 35 mm dishes with 1 μg of the WT or phospho-mutant Na_V1_.5 plasmid and/or 1.5 μg of the iFGF13A plasmid using 2 μL of Lipofectamine 2000 (Life technologies) following the manufacturer’s instructions. The empty pcDNA3.1 plasmid (Life Technologies) was used as a filler plasmid to keep the total DNA constant at 2 μg in each transfection.

### Immunoprecipitations

Forty-four hours after transfection, HEK-293 cells were washed twice with PBS and lysed in ice-cold lysis buffer containing 20 mM HEPES (pH 7.4), 150 mM NaCl, 5% Lauryl Maltose Neopentyl Glycol (Anatrace), 0.5% Cholesterol Hemisuccinate Tris Salt (Anatrace), 1X complete protease inhibitor cocktail tablet (Roche), 1 mM phenylmethylsulfonyl fluoride (PMSF, Interchim), 0.7 μg/ml pepstatin A (Thermo Fisher Scientific) and 1X Halt phosphatase inhibitor cocktail (Thermo Fisher Scientific). Soluble protein fractions were collected and subsequently used for immunoprecipitations (IP) with 1 μg of an anti-Na_V1_.5 rabbit polyclonal antibody (RbαNa_V1_.5, Merck, #S0819), raised against an epitope located in the first intracellular linker loop of Na_V1_.5. Prior to the IP, antibodies were cross-linked to protein G-magnetic beads using 20 mM dimethyl pimelimidate (Thermo Fisher Scientific) (80). Protein samples and antibody-coupled beads were mixed for 2 hours at 4°C. Magnetic beads were then collected, washed rapidly three times with ice-cold lysis buffer, and isolated protein complexes were eluted with 1X SDS sample buffer at 60°C for 10 min.

Total lysates and immunoprecipitated proteins were fractionated on one-dimensional polyacrylamide gels and analyzed by western blotting using a rabbit monoclonal anti-Na_V1_.5 antibody (1:2000, Cell Signaling, #14421) and a mouse monoclonal anti-PAN-FHF-A antibody (1:300, UC Davis/NINDS/NIMH NeuroMab Facility, clone N235/22) as described previously (80). Bound antibodies were detected using horseradish peroxidase-conjugated goat anti-rabbit or anti-mouse secondary antibodies (Santa Cruz Biotechnology), and protein signals were visualized using the SuperSignal West Dura Extended Duration Substrate (Pierce). The intensities of co-immunoprecipitated iFGF13A bands were normalized to the intensities of immunoprecipitated Na_V1_.5 bands from the same IP sample, and iFGF13A abundances from immunoprecipitations of Na_V1_.5-Y1495E channels are expressed relative to abundances in immunoprecipitations of Na_V1_.5-WT channels.

## Data availability

TEVC and VCF data along with simulation trajectories will be made available on Zenodo repository.

## ACKNOWLEDGMENTS

This project has received financial support by Lundbeck Foundation (R383-2022-165) and by the European Union’s Horizon 2020 research and innovation program under the Marie Skłodowska-Curie grant agreement No. 101109359. The computational work was supported by the Australian Research Council (DP250100893) and with the assistance of resources from the National Computational Infrastructure (NCI Australia), an NCRIS-enabled capability supported by the Australian Government. The authors would like to thank Dr. Hendrik Harms and Dr. Michelle Nilsson for the critical reading of the manuscript.

