## Supplementary Information for "Phosphorylation elicits bifurcated regulation of Na_V_1.5 by intracellular auxiliary proteins"

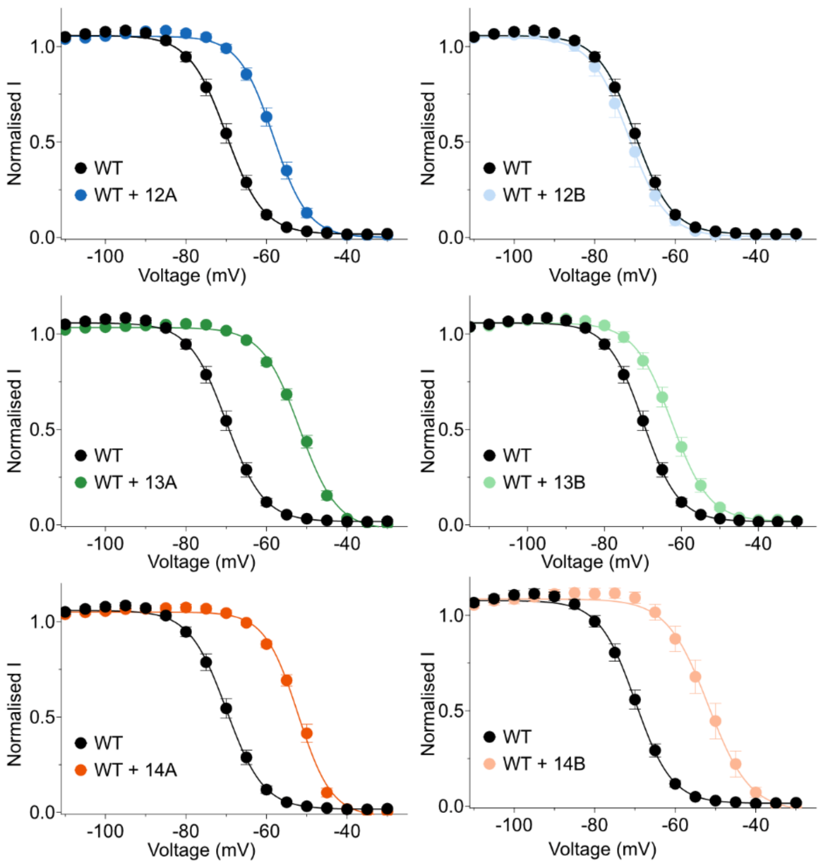


**Supplementary Figure S1**. **Effects of iFGFs on WT hNav1.5.** SSI curves for full-length hNav1.5 WT co-expressed with the different iFGFs isoforms investigated in this study. (mean +/-SEM, n=6-10)

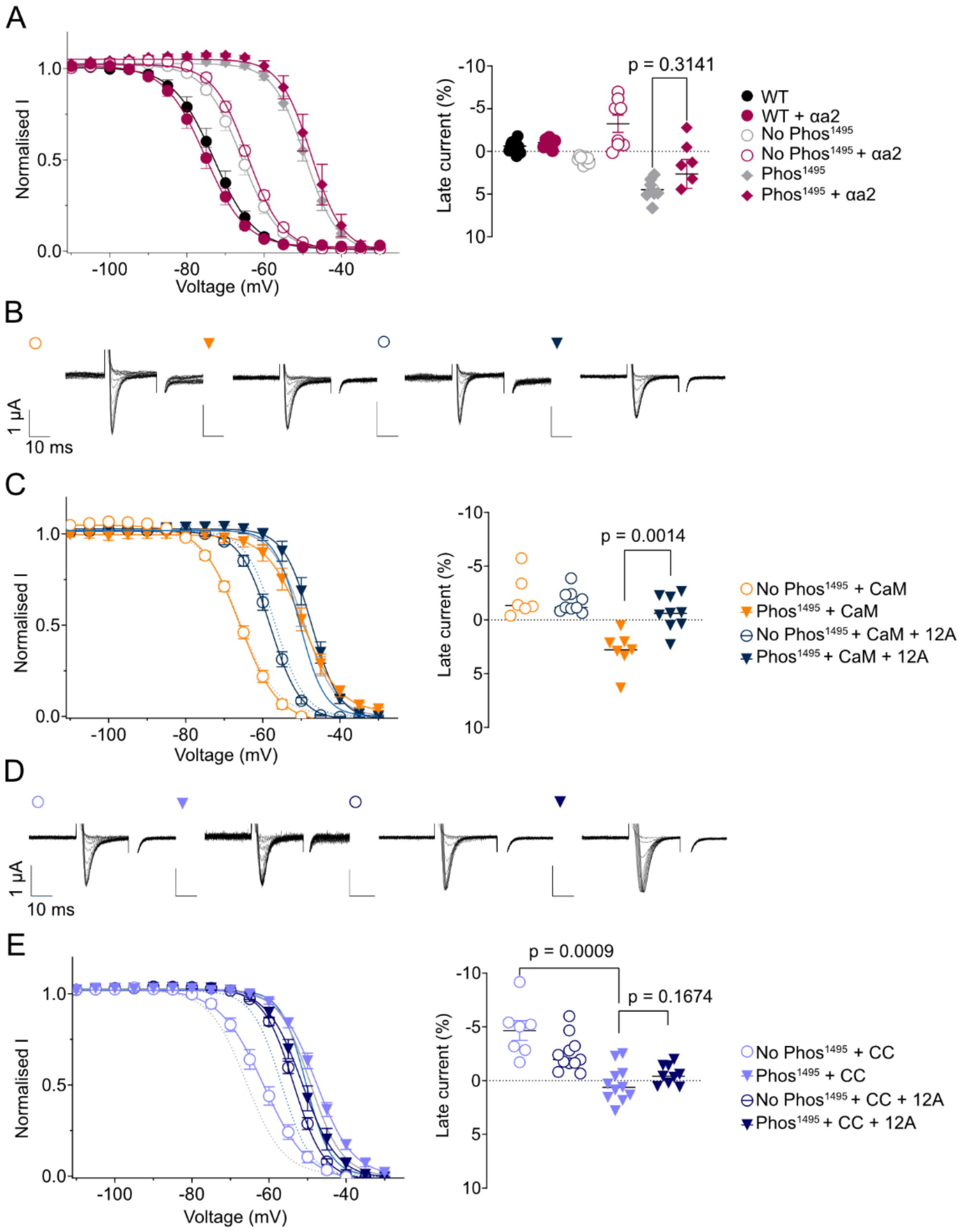


**Supplementary Figure S2: Alpha-actinin 2 (αα2) and calmodulin (CaM) overexpression or chelation does not impact iFGF12A effects. A.** SSI curves and late current properties of constructs No Phos$^{1495}$ and Phos$^{1495}$ co-expressed with αα2 and iFGF12A**. B.** Representative recordings of constructs No Phos$^{1495}$ and Phos$^{1495}$ co-expressed with CaM and iFGF12A. **C.** SSI curves and late current properties of the construct combinations shown in B. **D.** Representative recordings of constructs No Phos$^{1495}$ and Phos$^{1495}$ expressed with calmodulin chelator (CC) and iFGF12A. E. SSI curves and late current properties of the construct combinations shown in D. (mean +/-SEM, Welch's t-tests, n=6-20)

A

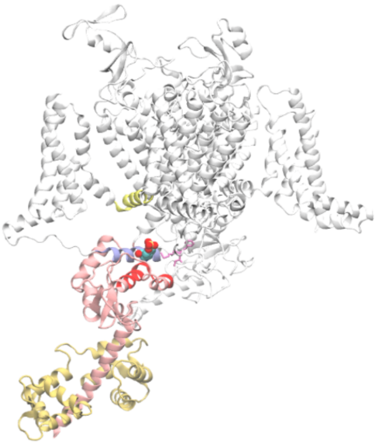

B

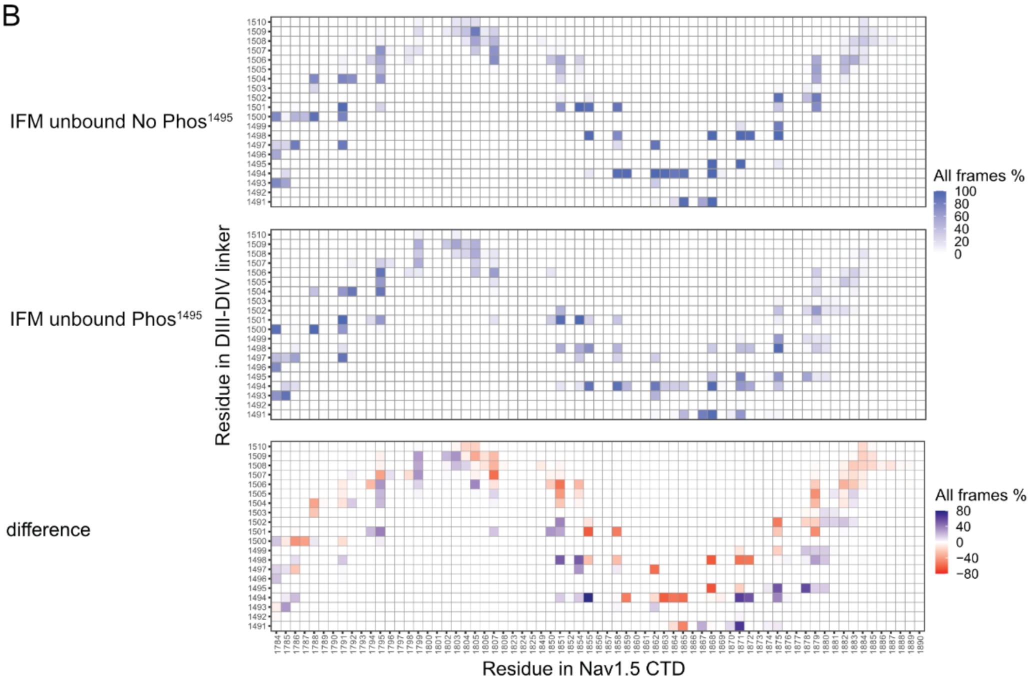
IFM unbound No Phos1495
IFM unbound Phos1495
difference
Residue in DIII-DIV linker
Residue in Nav1.5 CTD
All frames %
100
80
60
40
20
0
All frames %
80
40
0
−40
−80

C

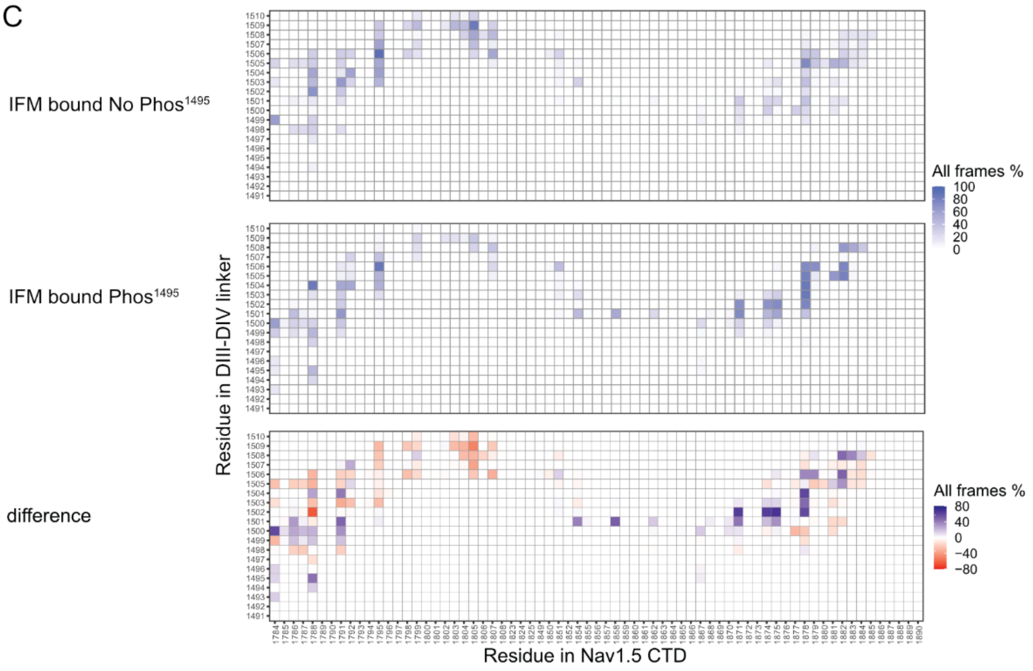
IFM bound No Phos1495
IFM bound Phos1495
difference
Residue in DIII-DIV linker
Residue in Nav1.5 CTD
All frames %
100
80
60
40
20
0
All frames %
80
40
0
−40
−80

**Supplementary Figure S3. Contact frequency between the residues in DIII-DIV linker and CTD of WT and Phos[1495] $Na_v$1.5 in IFM unbound and bound conformational states. A.** Model of the full length IFM-unbound state. DIV S4-S5 linker is shown in yellow, DIII-DIV linker in blue, CTD in pink, CaM in light brown, Y1495 in balls and the IFM motif in pink sticks. Proportion of frames residues from the DIII-DIV linker are in contact with those in the CTD is shown for the unphosphorylated (No Phos[1495] - top) and phosphorylated (Phos[1495] - middle) conditions and their difference for the (**B)** IFM-unbound (open) and (**C)** IFM-bound (inactivated) structures. A 5.0 Å distance cut-off was applied to determine whether a residue pair was in contact.

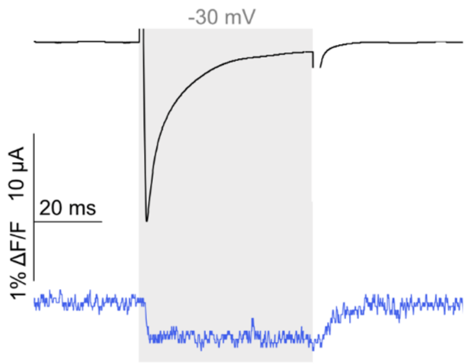


**Supplementary Figure S4. Voltage-clamp fluorometry recordings at Y1495 reveal a marked change of environment upon depolarization.** Representative VCF recording (current in black and fluorescence in blue) of Y1495-ANAP hNav1.5 in response to a depolarizing step to -30 mV (n=5).

```
C_1.5   SKKPQKPIPRPLNKYQGFIFDIVTKQAFDVTIMFLICLNMVTMMVETDDQSPEKINILAK   60
C_1.4   SKKPQKPIPRPQNKIQGMVYDLVTKQAFDITIMILICLNMVTMMVETDNQSQLKVDILYN   60

C_1.5   INLLFVAIFTGECIVKLAALRHYYFTNSWNIFDFVVVILSIVGTVLSDIIQKYFFSPTLF   120
C_1.4   INMIFIIIFTGECVLKMLALRQYYFTVGWNIFDFVVVILSIVGLALSDLIQKYFVSPTLF   120

C_1.5   RVIRLARIGRILRLIRGAKGIRTLLFALMMSLPALFNIGLLLFLVMFIYSIFGMANFAYV   180
C_1.4   RVIRLARIGRVLRLIRGAKGIRTLLFALMMSLPALFNIGLLLFLVMFIYSIFGMSNFAYV   180

C_1.5   KWEAGIDDMFNFQTFANSMLCLFQITTSAGWDGLLSPILNTGPPYCDPTLPNSN-GSRGD   239
C_1.4   KKESGIDDMFNFETFGNSIICLFEITTSAGWDGLLNPILNSGPPDCDPNLENPGTSVKGD   240

C_1.5   CGSPAVGILFFTTYIIISFLIVVNMYIAIILENFSVATEESTEPLSEDDFDMFYEIWEKF   299
C_1.4   CGNPSIGICFFCSYIIISFLIVVNMYIAIILENFNVATEESSEPLGEDDFEMFYETWEKF   300

C_1.5   DPEATQFIEYSVLSDFADALSEPLRIAKPNQISLINMDLPMVSGDRIHCMDILFAFTKRV   359
C_1.4   DPDATQFIAYSRLSDFVDTLQEPLRIAKPNKIKLITLDLPMVPGDKIHCLDILFALTKEV   360

C_1.5   LGESGEMDALKIQMEEKFMAANPSKISYEPITTTLRRKHEEVSAMVIQRAFRRHLLQRSL   419
C_1.4   LGDSGEMDALKQTMEEKFMAANPSKVSYEPITTTLKRKHEEVCAIKIQRAYRRHLLQRSM   420

C_1.5   KHASFLFRQQAGSGLSEEDAPEREGLIAYVMSENFSRPLGPPSSSSISSTSFPPSYDSVT   479
C_1.4   KQASYMYRHSHD--GSGDDAPEKEGLLANTMSKMYGHENGNSSSPSPEEKGEAG------   472

C_1.5   RATSDNLQVRGSDYSHSEDLADFPPSPDRDRE-------SIV                     514
C_1.4   -DAGPTM---G-LMPISPSDTAWPPAPPPGQTVRPGVKESLV                     509
```

**Supplementary Figure S5. C-terminal alignment of hNav1.5 and hNav1.4.** The alignment has been performed *via* Clustal Omega, Multiple Sequence Alignment (MSA) (EMBL-EBI (1)) and is represented without the split intein sequence that is similar between the two C constructs. Identical side chains are colored blue.

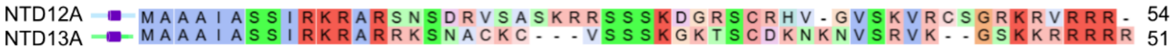


**Supplementary Figure S6.** NTD12A and NTD13A construct sequences alignment performed *via* Clustal Omega, Multiple Sequence Alignment (MSA) (EMBL-EBI (1)) colored by the chemical nature and conservation level of the residues.

**Supplementary Table 1. $V_{50}$SSI, $V_{50}$Act and late current properties for the constructs investigated in this study.** All data are from at least two batches of oocytes. For values marked with * the exclusion criteria have been altered due to small currents (currents exceeding -0.2 μA, versus -0.5 μA for all the others, see also Methods). Late currents greater than 2% are highlighted in italic font.

| Construct | $V_{50}$SSi +/- SEM (n) (mV) | $I_{late}$ +/- SEM (n) (%) | $V_{50}$Act +/- SEM (n) (mV) |
|---|---|---|---|
| Wild type hNa$_V$1.5 (WT) | -69.881 +/- 0.270 (8) | -1.319 +/- 0.243 (20) | -34.562 +/- 0.569 (16) |
| E1784K | -74.799 +/- 0.470 (9) | *4.216 +/- 0.583 (8)* | -27.349 +/- 1.572 (6) |
| E1784K + iFGF12A | -69.882 +/- 0.242 (13) | -1.664 +/- 0.216 (8) | -35.581 +/- 0.508 (7) |
| E1784K + iFGF13A | -64.501 +/- 0.315 (12) | -1.451 +/- 0.309 (12) | -34.817 +/- 0.852 (7) |
| WT + iFGF12A | -58.265 +/- 0.270 (6) | -2.574 +/-0.735 (6) | -38.587 +/- 1.315 (6) |
| WT + iFGF12B | -71.496 +/- 0.432 (12) | -2.194 +/- 0.497 (12) | -38.182 +/- 0.687 (9) |
| WT + iFGF13A | -51.764 +/- 0.234 (8) | -0.688 +/- 0.201 (7) | -33.751 +/- 0.724 (5) |
| WT + iFGF13B | -62.513 +/- 0.333 (10) | -2.697 +/- 0.468 (10) | -35.200 +/- 0.564 (10) |
| WT + iFGF14A | -52.149 +/- 0.261 (6) | -1.750 +/- 0.317 (11) | -34.353 +/- 0.468 (8) |
| WT + iFGF14B | -51.819 +/- 0.684 (8) | -1.916 +/- 0.229 (10) | -28.344 +/- 0.824 (6) |
| WT + NtFGF12A | -66.448 +/- 0.421 (10) | -0.467 +/- 0.330 (5) | -25.437 +/- 1.269 (8) |
| WT + NtFGF13A | -56.187 +/- 0.563 (8) | -0.082 +/- 0.386 (7) | -30.100 +/- 0.293 (6) |
| WT + α-actinin-2 | -75.275 +/- 0.364 (9) | -1.044 +/- 0.2047 (7) | -43.196 +/- 0.499 (5) |
| No Phos$^{1495}$ | -65.849 +/- 0.294 (8) | -1.262 +/- 0.374 (14) | -38.068 +/ 0.430 (7) |
| Phos$^{1495}$ | -49.322 +/- 0.329 (7) | *3.287 +/- 0.262 (13)* | -36.960 +/- 0.419 (5) |
| No Phos$^{1495}$ + iFGF12A | -56.795 +/- 0.357 (7) | -1.933 +/- 0.256 (7) | -38.279 +/- 0.371 (5) |
| Phos$^{1495}$ + iFGF12A | -50.419 +/- 0.197 (12) | -0.971 +/- 0.392 (9) | -39.283 +/- 0.661 (7) |
| No Phos$^{1495}$ + iFGF12B | -63.831 +/- 1.196 (6) | -1.916 +/- 0.776 (7) | -39.892 +/- 0.672 (8) |
| Phos$^{1495}$ + iFGF12B | -55.546 +/- 1.416 (5) | 1.215 +/- 0.495 (7) | -42.551 +/- 0.913 (6) |
| No Phos$^{1495}$ + iFGF13A | -49.017 +/- 0.744 (5) | -1.368 +/- 0.316 (5) | -32.420 +/- 1.054 (5) |
| Phos$^{1495}$ + iFGF13A | -47.236 +/- 0.684 (5) | 0.506 +/- 0.510 (7) | -38.196 +/- 1.833 (7) |
| No Phos$^{1495}$ + iFGF13B | -60.887 +/- 0.770 (7) | -2.049 +/- 0.672 (7) | -33.522 +/- 2.909 (5) |
| Phos$^{1495}$ + iFGF13B | -44.102 +/- 1.088 (5) | *2.967 +/- 0.279 (6)* | -29.621 +/- 1.266 (6) |
| No Phos$^{1495}$ + iFGF14A | -58.613 +/- 0.699 (6) | -0.717 +/- 0.132 (9) | -37.137 +/- 1.690 (5) |
| Phos$^{1495}$ + iFGF14A | -50.644 +/- 0.725 (5) | -0.015 +/- 0.211 (6) | -38.275 +/- 1.173 (5) |
| No Phos$^{1495}$ + iFGF14B | -53.902 +/- 1.087 (7) | -1.791 +/- 0.687 (7) | -30.150 +/- 1.123 (6) |
| Phos$^{1495}$ + iFGF14B | -45.511 +/- 1.868 (8) | *3.433 +/- 0.742 (6)* | -24.528 +/- 0.949 (8) |
| No Phos$^{1495}$ + NTD12A | -58.918 +/- 0.351 (10) | -2.097 +/- 0.733 (9) | -30.112 +/- 0.360 (9) |
| Phos$^{1495}$ + NTD12A | -48.910 +/- 0.640 (8) | 0.812 +/- 0.473 (8) | -30.749 +/- 0.452 (8) |
| No Phos$^{1495}$ + NTD13A | -57.828 +/- 0.435 (8) | 0.093 +/- 0.138 (9) | -35.717 +/- 0.738 (5) |
| Phos$^{1495}$ + NTD13A | -47.236 +/- 0.684 (5) | *3.332 +/- 0.300 (8)* | -37.384 +/- 0.891 (7) |
| E1784K No Phos$^{1495}$ (*) | -60.330 +/- 0.595 (7) | -1.682 +/- 0.880 (6) | -28.036 +/- 0.416 (5) |
| E1784K Phos$^{1495}$ | -52.017 +/- 0.459 (7) | *4.285 +/- 0.719 (9)* | -31.257 +/- 1.321 (8) |
| E1784K No Phos$^{1495}$ + FGF12A | -61.131 +/- 0.526 (8) | -4.837 +/- 1.015 (6) | -30.393 +/- 0.407 (8) |
| E1784K Phos$^{1495}$ + iFGF12A | -48.339 +/- 0.253 (10) | -1.745 +/- 0.500 (5) | -28.416 +/- 0.421 (6) |
| E1784K No Phos$^{1495}$ + iFGF13A | -59.284 +/- 0.388 (6) | -3.916 +/- 0.823 (6) | -32.300 +/- 0.688 (6) |
| E1784K Phos$^{1495}$ + iFGF13A | -47.406 +/- 0.877 (12) | -0.538 +/- 0.480 (10) | -34.209 +/- 0.885 (6) |
| No Phos$^{1495}$ + α-actinin-2 | -64.203 +/- 0.269 (8) | -3.237 +/- 1.003 (7) | -38.169 +/- 0.719 (6) |
| Phos$^{1495}$ + α-actinin-2 | -47.745 +/- 0.592 (7) | *2.642 +/- 1.686 (7)* | -36.785 +/- 0.499 (6) |

**Nav1.5-Nav1.4 chimera experiment**

| | | | |
|---|---|---|---|
| Nav15-Nav14 No Phos$^{1495}$ | -63.849 +/- 0.497 (12) | -3.954 +/- 1.109 (5) | -30.791 +/- 0.991 (7) |
| Nav15/Nav14 Phos$^{1495}$ | -45.057 +/- 0.507 (10) | *3.530 +/- 0.722 (15)* | -31.375 +/- 0.518 (7) |

| | | | |
|---|---|---|---|
| Nav15-Nav14 No Phos$^{1495}$ + iFGF12A | -52.895 +/- 0.414 (9) | -1.585 +/- 0.641 (6) | -29.261 +/- 0.783 (9) |
| Nav15/Nav14 Phos$^{1495}$ + iFGF12A | -42.617 +/- 0.576 (9) | 0.883 +/- 0.397 (8) | -30.077 +/- 0.586 (8) |

**Calmodulin & calmodulin chelator experiment**

| | | | |
|---|---|---|---|
| CaM No Phos$^{1495}$ | -66.007 +/- 0.256 (9) | -2.210 +/- 0.815 (6) | -37.789 +/- 0.210 (8) |
| CaM Phos$^{1495}$ | -49.646 +/- 0.539 (9) | *2.865 +/- 0.671 (7)* | -35.458 +/- 0.738 (7) |
| CaM + 12A No Phos$^{1495}$ | -58.102 +/- 0.215 (10) | -1.711 +/- 0.344 (9) | -37.374 +/- 0.273 (7) |
| CaM + 12A Phos$^{1495}$ | -47.361 +/- 0.393 (9) | -0.626 +/- 0.525 (9) | -34.334 +/- 0.406 (7) |
| CC No Phos$^{1495}$ | -61.375 +/- 0.406 (13) | -4.656 +/- 0.920 (7) | -31.173 +/- 0.532 (6) |
| CC Phos$^{1495}$ | -47.736 +/- 0.328 (14) | 0.391 +/- 0.502 (11) | -27.242 +/- 0.757 (4) |
| CC 12a No Phos$^{1495}$ | -53.150 +/- 0.691 (22) | -2.507 +/- 0.487 (11) | -30.492 +/- 0.494 (5) |
| CC 12a Phos$^{1495}$ | -53.577 +/- 0.181 (17) | -0.529 +/- 0.304 (9) | -40.091 +/- 0.529 (4) |
| WT + CC | -70.284 +/- 0.366 (9) | -3.478 +/- 0.454 (12) | -30.951 +/- 0.396 (6) |
| WT + CC + 12A | -60.984 +/- 0.354 (9) | -1.699 +/- 0.299 (7) | -32.596 +/- 0.321 (5) |
| WT + CaM | -70.110 +/- 0.164 (10) | -2.650 +/- 0.498 (13) | -30.976 +/- 0.321 (8) |
| WT + CaM + 12A | -62.457 +/- 0.268 (6) | -2.964 +/- 0.671 (7) | -31.462 +/- 0.342 (7) |
